# Differential clearance rate of proteins encoded on a self-amplifying mRNA vaccine in muscle and lymph nodes

**DOI:** 10.1101/2024.10.07.615730

**Authors:** Reo Kanechi, Tatsuya Shishido, Mana Tachikawa, Tomohiro Nishimura, Akihito Sawada, Hayato Okade, Daisuke Ishikawa, Hitoshi Yamaguchi, Marito Araki

## Abstract

ARCT-154, a recently approved self-amplifying mRNA (saRNA) vaccine, has been shown in clinical trials to induce higher levels of neutralizing antibodies and sustain them for a longer time than the conventional mRNA vaccine BNT162b2. However, the scientific evidence explaining this superiority remain elusive. Hence, we explored the temporal changes in spike protein and replicase components following a single dose of ARCT-154 vaccination in mice. The encoded spike protein reached its highest level approximately 3 days after vaccination and quickly disappeared from the rectus femoris muscle, the injection site. Although the spike protein levels also peaked at an early time point in the lymph nodes, it remained detectable 28 days after the vaccination and then disappeared by 44 days after the vaccination. Expression of nsP1, nsP2 and nsP4 was observed in the injected muscle and/or the lymph nodes for up to 15 days post-vaccination. These data suggest that prolonged expression of spike proteins in lymph nodes may, if not entirely, be responsible for the induction of higher and prolonged levels of neutralizing antibodies by the saRNA vaccine.

## Introduction

The coronavirus disease 2019 (COVID-19) pandemic has resulted in the deaths of more than 7 million people. During this time, we saw the rapid development and use of effective vaccines against severe acute respiratory syndrome coronavirus 2 (SARS-CoV-2) [1,2]. Although the mRNA vaccine against SARS-CoV-2 showed high efficacy against the original strain, the duration of immunity induced by the mRNA was relatively short; the situation was further exacerbated by immune evasion by mutant strains, resulting in reduced efficacy[3,4]. With the ongoing public health burden of SARS-CoV-2, first generation vaccines are sub-optimal for continued ongoing use as we transition out of the pandemic. Most notably, there remains a need for a more durable vaccine. To develop novel vaccines that induce long-lasting immune responses and are highly cross-reactive to a broad range of variants, it is essential to deepen our understanding of the mechanisms of action of mRNA vaccines.

ARCT-154, the world’s first approved self-amplifying mRNA (saRNA) vaccine originally developed by Arcturus Therapeutics (San Diego, CA, USA), consists of mRNA formulated with lipid nanoparticles (LNP) harboring the Venezuelan equine encephalitis virus (VEEV) genome, which has been modified by replacing the structural genes essential for virus infection with the gene encoding the D614G variant form of the SARS-CoV-2 spike protein[5-7]. This modification eliminates infectivity and allows for the expression of the spike protein. In a phase 3 clinical trial, a booster dose of ARCT-154 induced noninferior immunogenicity 1 month after administration and production of longer-lasting neutralizing antibodies compared with the conventional mRNA vaccine BNT162b2[5,6]. Following the confirmation of safety profiles in clinical trials[5], ARCT-154 was approved in Japan as the first saRNA vaccine in the world[8]. These data suggest that saRNA vaccines are likely to exhibit immunogenicity that induces persistent immune responses. However, the mechanisms underlying the long-lasting immune response induced by saRNAs have yet to be fully explored.

In the mRNA of ARCT-154, two open reading frames (ORFs) encode a four-subunit VEEV replicase that consists of non-structural proteins (nsP)1–4 and the SARS-CoV-2 spike protein. In the target cells, the replicase synthesizes the complementary negative-sense strand of the saRNA, which is later used as a template by the replicase to generate a complementary positive-sense saRNA in a self-amplification process. The replicase also synthesizes a small subgenomic positive-sense RNA containing the second ORF, which is translated by the host cells to produce the SARS-CoV-2 spike protein, by recognizing the subgenomic promoter in the negative saRNA strand[9,10]. In this way, the mRNA contained in the saRNA vaccine is amplified in host cells, resulting in increased and prolonged antigen protein expression in mice administrated with the saRNA vaccine than the protein expression in mice administrated an equivalent dose of conventional mRNA vaccine[11].

After vaccination, it is expected that the saRNA vaccine would be degraded and eliminated by the body in a manner similar to that of conventional mRNA vaccines. However, the pharmacokinetics of the protein encoded by a clinically approved saRNA vaccine are unknown. In this study, we used mice to determine the biodistribution of the SARS-CoV-2 spike protein and replicase components encoded by the saRNA vaccine. We examined the temporal changes in these proteins in tissues, including the injection site (rectus femoris muscle), spleen, lymph nodes, and serum, after a single-dose ARCT-154 vaccination.

## Materials and methods

### Self-Amplifying RNA vaccine

ARCT-154 was obtained from Arcturus Therapeutics. The vaccine was supplied in a vial containing 100 µg active ingredient and was stored at -20°C or lower before use. It was dissolved and diluted in sterile saline for use in the immunization experiments.

### Immunization

Female BALB/c mice aged 8 weeks (Japan SLC) were intramuscularly immunized through the rectus femoris with ARCT-154 at a dose of 10 µg in 50 µL of saline (n = 5 mice/interval). Mice were sacrificed at 5 hours and 1, 2, 3, 5, 7, 15, 28, and 44 days after vaccination, and the rectus femoris muscles, inguinal lymph nodes, spleens, and serum were harvested for analysis. The study protocol was approved by the KM Biologics Institutional Animal Care and Use Committee (#R240612-2B).

### Tissue lysate preparation

Tissue samples were loaded into Lysing Matrix D 2mL tubes (MP Biomedicals) filled with radioimmunoprecipitation assay lysis buffer plus protease inhibitor cocktail (Santa Cruz Biotechnology) and homogenized using FastPrep-24 5G (MP Biomedicals). Tissue samples were sonicated for 45 sec, followed by a centrifugation step at 10,000 ×*g* for 10 min at 4°C to remove debris. Protein concentrations of tissue lysates were determined using bicinchoninic acid assay (Thermo Fisher Scientific), and lysates were stored at -80°C.

### Dot blot analysis

Tissue lysates were diluted with phosphate-buffered saline (PBS) to a final protein concentration of 1 µg/µL, and 2 µg protein was then spotted onto a nitrocellulose membrane (Bio-Rad). The membrane was air-dried and then blocked in Blocking One (NACALAI TESQUE) or 5% skim milk (FUJIFILM Wako chemicals) in PBS with 0.05% Tween 20 (PBS-T) at 4°C overnight. To detect the SARS-CoV-2 Spike protein, the membrane was incubated with an anti-S protein antibody (Abnova, clone: BHB1) diluted to a concentration of 1/5000 in Can Get Signal Solution 1 (TOYOBO) at room temperature for 1 hour. To detect VEEV nsP1, nsP2, nsP3, and nsP4, the membrane was incubated with anti-nsP1 antibody (GeneTex, clone HL1472), anti-nsP2 antibody (GeneTex, clone HL1919), anti-nsP3 antibody (GeneTex, clone HL1502), or anti-nsP4 antibody (GeneTex, clone HL1741), respectively, diluted to a concentration of 1/5,000 in 5% skim milk at room temperature for 1 hour. The membranes were washed twice with PBS-T for 10 min each and then incubated with horseradish peroxidase (HRP)-conjugated goat anti-human IgG (Abcam) diluted to a concentration of 1/5,000 in Can Get Signal Solution 2 (TOYOBO) or HRP-conjugated goat anti-rabbit IgG (Bio-Rad) diluted to a concentration of 1/3,000 in 5% skim milk at room temperature for 1 hour. After the membranes were washed twice with PBS-T for 10 min each, chemiluminescence was detected using Western BLoT Ultrasensitive HRP substrate (TaKaRa) and LuminoGraph I (ATTO). The signal density of the dots was quantified with densitometry using the Volume Tools of the CS Analyzer 4 software (ATTO). Densities obtained with the respective tissue in non-immunized mice were set as the baseline.

### Statistical analysis

Statistical analyses were performed using GraphPad Prism 10 software version 10.2.3 (GraphPad Software). Unpaired two-tailed Mann–Whitney U-tests were used to determine the significance of differences. *p* < 0.05 were considered statistically significant.

## Results

### Differential clearance rate of spike protein in tissues following ARCT-154 vaccination

To examine the tissue distribution and temporal changes of proteins encoded on ARCT-154 (Fig. 1A), 10 µg of ARCT-154 was intramuscularly injected into the left hind leg muscle of BALB/c mice, and lysates were prepared from tissues collected at each time point after the vaccination (Fig. 1B). As shown in Fig. 1C, SARS-CoV-2 spike protein was detected in the rectus femoris muscles, the site of injection, 5 hours after vaccination, and its levels peaked at around 3 days after vaccination; the levels of spike proteins sharply decreased 7 days after vaccination and became undetectable 15 days after vaccination or thereafter (Fig. 1C). However, in inguinal lymph nodes, the levels of spike protein peaked at around 1 day after vaccination, gradually decreased, and eventually dropped to baseline level 44 days after vaccination (Fig. 1D). Spike proteins were undetectable in the spleen (Fig. 1E) or serum (Fig. 1F) at all time points.

**Figure 1.**
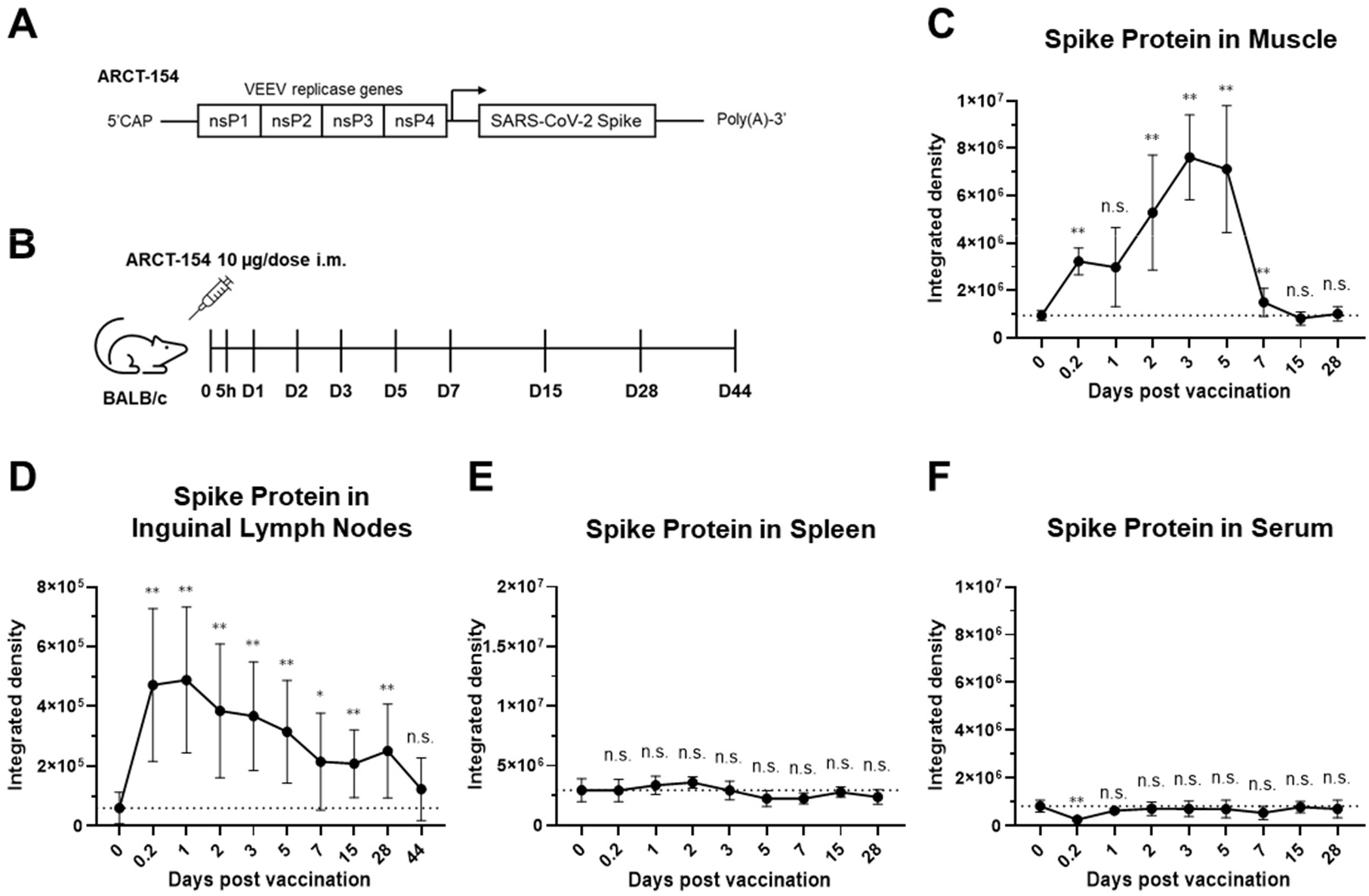
Temporal changes of spike protein expression in tissues following a single administration of the ARCT-154 vaccine. (A) Schematic illustration of the saRNA in ARCT-154 vaccine. Arrow indicates the subgenomic promoter. nsP, non-structural protein. VEEV, Venezuelan equine encephalitis virus. SARS-CoV-2, severe acute respiratory syndrome coronavirus 2. (B) Schematic representation of the experimental procedure. BALB/c mice (n = 5 mice/interval) were intramuscularly (i.m.) immunized with ARCT-154, and tissues were harvested at indicated time points. (C–F) Levels of SARS-CoV-2 spike protein in tissue lysates of rectus femoris muscles (C), inguinal lymph nodes (D), spleen (E), and serum (F) of mice at different intervals after vaccination with ARCT-154 determined by dot blot analysis. Values are shown as mean with a 95% confidence interval. Signal intensities obtained with the respective tissue in non-immunized mice were set as the baseline, indicated by the dotted line. Significance was determined using unpaired two-tailed Mann–Whitney U-tests. **p < 0.01. *p < 0.05. n.s., not significant, compared to background (time 0).

### Clearance of nsP1 in tissues following ARCT-154 vaccination

Since nsP1 is expected to be translated first among all genes encoded on ARCT-154, we analyzed the temporal changes in nsP1 in animals. In the rectus femoris muscle, nsP1 was detected from 2 days after vaccination and peaked at around 3 days after vaccination; the levels of nsP1 gradually decreased and reached background levels 28 days after vaccination (Fig. 2A). In the inguinal lymph nodes, nsP1 protein was detected starting from 5 hours after vaccination and peaked by 1 day after vaccination. The levels of nsP1 declined and became undetectable by 15 days after vaccination (Fig. 2B). nsP1 was undetectable in the spleen at all time points (Fig. 2C).

**Figure 2.**
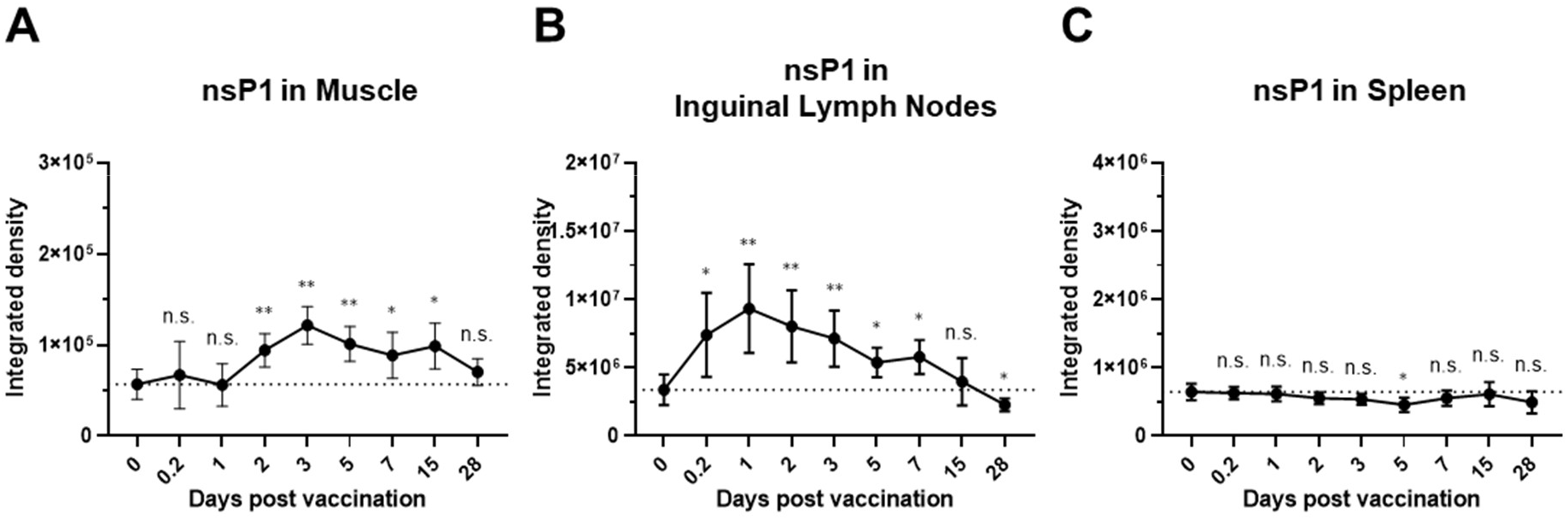
Temporal changes of nsP1 in tissue following a single administration of the ARCT-154 vaccine. (A–C) Levels of nsP1 in tissue lysates of rectus femoris muscles (A), inguinal lymph nodes (B), and spleen (C) of mice at different intervals after vaccination with ARCT-154 determined by dot blot analysis. Values are shown as mean with a 95% confidence interval. Signal intensities obtained with the respective tissue in non-immunized mice were set as the baseline, indicated by the dotted line. Significance was determined using unpaired two-tailed Mann–Whitney U-tests. **p < 0.01. *p < 0.05. n.s., not significant, compared to background (time 0). nsP, non-structural protein.

### Clearance of nsP2–4 in tissues following ARCT-154 vaccination

We examined the expression of nsP2–4 encoded by ARCT-154 in the rectus femoris muscles and inguinal lymph nodes where nsP1 was detected (Fig. 2). Unlike nsP1, nsP2-specific signal was undetectable in muscles at all time points, presumably due to the higher background signal of detecting antibody for the lysate (Fig. 3A). nsP3 was barely detectable in the muscle between 2 and 3 days after vaccination (Fig. 3B). nsP4 in the rectus femoris muscle gradually accumulated and became undetectable by 15 days after vaccination (Fig. 3C). In the inguinal lymph nodes, nsP2 expression profile resembled to spike protein. nsP2 was detected 5 hours after vaccination, persisted for some time, and eventually became undetectable by 28 days after vaccination (Fig. 3D). nsP3 was undetectable at all time points (Fig. 3E). nsP4 was detected 5 hours after vaccination and decreased to background levels by 15 days after vaccination (Fig. 3F).

**Figure 3.**
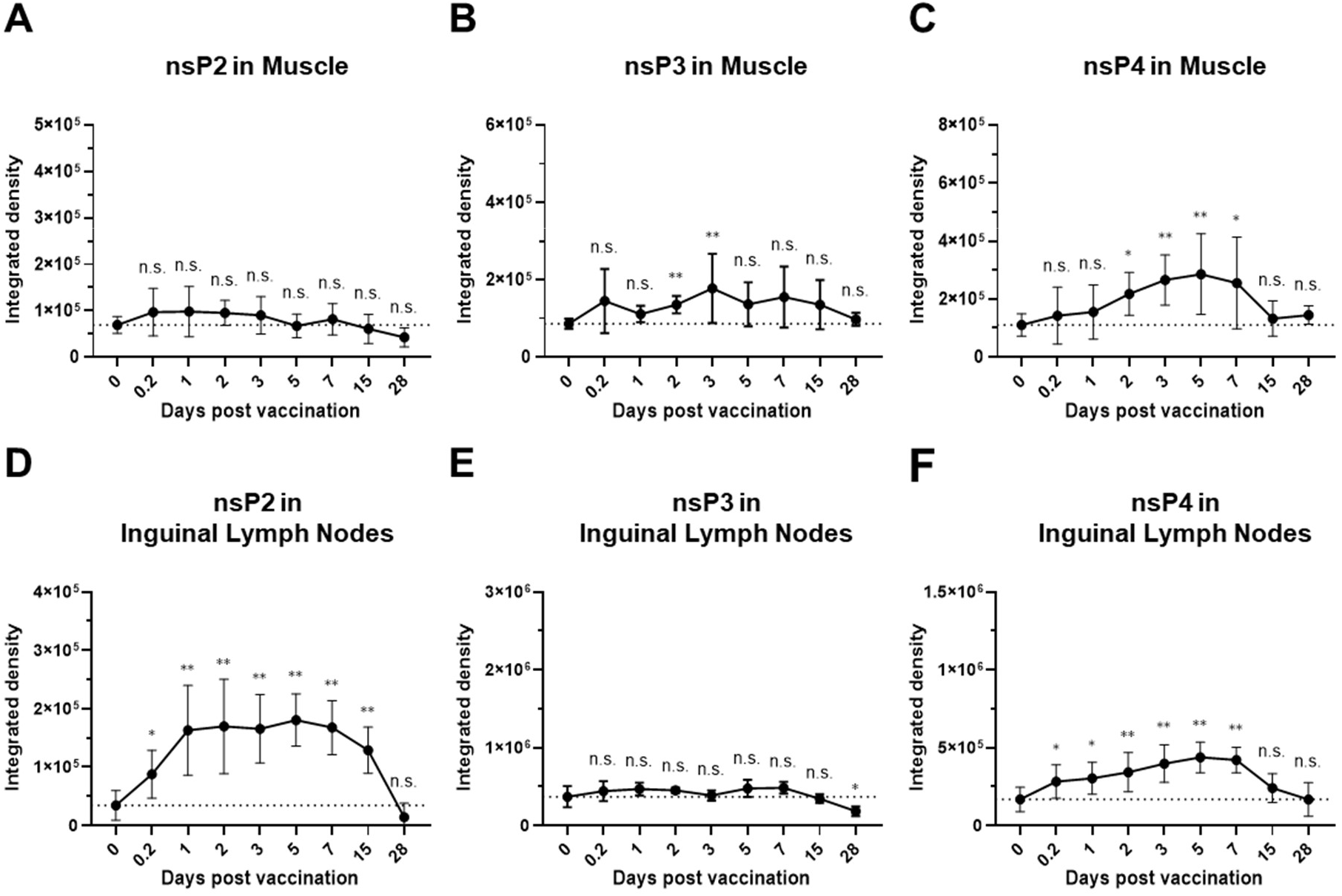
Temporal changes of nsP2–4 in rectus femoris muscle and inguinal lymph nodes following a single administration of ARCT-154 vaccine. (A–F) Levels of nsP2 (A, D), nsP3 (B, E), and nsP4 (C, F) in tissue lysates of rectus femoris muscles (A–C) and inguinal lymph nodes (D–F) of mice at different intervals after vaccination with ARCT-154 determined by dot blot analysis. Values are shown as mean with a 95% confidence interval. Signal intensities obtained with the respective tissue in non-immunized mice were set as the baseline, indicated by the dotted line. Significance was determined using unpaired two-tailed Mann–Whitney U-tests. **p < 0.01. *p < 0.05. n.s., not significant, compared to background (time 0). nsP, non-structural protein.

## Discussion

We showed temporal changes in spike protein and replicase components following vaccination with a single intramuscular dose of ARCT-154. Spike protein levels peaked approximately 3 days after vaccination and rapidly disappeared at the injection site (rectus femoris muscle) (Fig. 1C). Although it is difficult to draw conclusions owing to the challenges of detecting specific signals in muscle tissue, nsP1 and nsP4 expression seemed to mirror that of the spike protein (Fig. 2A, Fig. 3C). In the lymph nodes, the spike protein peaked at an early time point and gradually decreased, becoming undetectable by 44 days after vaccination (Fig. 1D). A gradual decrease of nsP2 in the lymph nodes was also observed (Fig. 3D). These data suggest that prolonged expression of spike proteins in lymph nodes may, if not entirely, be responsible for the induction of higher and prolonged levels of neutralizing antibodies by the saRNA vaccine.

Previous studies investigating temporal changes in proteins encoded by the mRNA of saRNA vaccines have reported that the disappearance of the protein occurs by 28[12] or 63[13] days after vaccination. Despite the difference in experimental systems and conditions, the period of persistent expression of the encoded protein on ARCT-154 (28 days or more but less than 44 days) was comparable to that in previous reports. Contrastingly, a shorter persistence of the encoded protein, such as 7 days[14], was observed with the conventional mRNA vaccine. However, a direct comparison of saRNA and conventional mRNA vaccines is required to determine differences in the tissue distribution of target proteins.

Although the vaccine was injected to rectus femoris muscle, proteins encoded on saRNA were detected in the lymph nodes, even at early time points (Fig. 1D, 2B, 3D, 3F). A recent study using positron emission tomography-tracer-labeled LNP revealed a rapid transition of LNP to the lymph nodes from the injection site[15], which is consistent with our observations.

The rapid clearance of spike protein from the rectus femoris muscle may be owing to adaptive immune response against the spike protein. In lymph nodes, immune cells may be less susceptible to attacks from other immune cells, allowing them to maintain prolonged expression of encoded proteins. Detailed analyses of the persistence of target antigen expression and mRNA functionality in lymph nodes, more specifically, in antigen-presenting cells after the administration of saRNA vaccines versus traditional vaccines are necessary to further understand the reasons for the longer duration and higher levels of neutralizing antibody titers induced by saRNA vaccines.

Limitations of this study include a lack of quantitative determination of protein concentration, suboptimal detection of replicase components (e.g. nsP3), and a limited variety of organs studied. More comprehensive studies that include various organs, using highly sensitive and quantitative measurements of encoded proteins, are required to gain a more detailed understanding of the in vivo distribution and dynamics of proteins expressed by saRNA vaccines.

In conclusion, we observed transient expression of the spike protein at the injection site and a relatively prolonged expression in lymph nodes, which might explain the strong and sustained neutralizing antibody response induced by the saRNA vaccine. Comparing the differences between saRNA and conventional mRNA vaccines will provide insight into the mechanisms behind the enhanced immune response observed with saRNA vaccines in clinical trials, contributing to the development of safer and more effective mRNA vaccines.

## Author contributions: CRediT

**Reo Kanechi:** Conceptualization, Data curation, Formal analysis, Investigation, Methodology, Writing– review & editing. **Tatsuya Shishido:** Conceptualization, Data curation, Formal analysis, Investigation, Methodology, Project administration, Writing – original draft, Writing – review & editing. **Mana Tachikawa:** Data curation, Formal analysis, Writing – review & editing. **Tomohiro Nishimura:** Conceptualization, Investigation, Methodology, Writing – review and editing. **Akihito Sawada:** Investigation, Writing – review & editing. **Hayato Okade:** Investigation, Writing – review & editing. **Daisuke Ishikawa:** Investigation, Writing – review & editing. **Hitoshi Yamaguchi:** Conceptualization, Writing – review & editing. **Marito Araki:** Conceptualization, Investigation, Methodology, Supervision, Writing – original draft, Writing – review & editing.

## Funding

This study was conducted without financial support from any external organizations.

## Declaration of competing interest

Reo Kanechi, Akihito Sawada, Hayato Okade, Hitoshi Yamaguchi, and Marito Araki are employees of Meiji Seika Pharma Co. Ltd. Tatsuya Shishido, Mana Tachikawa, Tomohiro Nishimura, Daisuke Ishikawa, Hitoshi Yamaguchi, and Marito Araki are employees of KM Biologics Co. Ltd.

## Acknowledgments

We are grateful to Sachiko Kuriyama, Ruriko Miyamoto, Yuka Kanadome, and Tomoyo Tsuneyoshi for technical assistance. We thank Arcturus Therapeutics, Inc., and CSL Seqirus for their critical review of the manuscript.

## Abbreviations

saRNA: self-amplifying mRNA
nsP: non-structural protein
COVID-19: coronavirus disease 2019
SARS-CoV-2: severe acute respiratory syndrome coronavirus 2
VEEV: Venezuelan equine encephalitis virus
ORF: open reading frames
PBS: phosphate-buffered saline
PBS-T: phosphate-buffered saline with Tween 20
HRP: horseradish peroxidase
LNP: lipid nanoparticles

## References

[1] J. Kates, J. Michaud, Ten Numbers to Mark Three Years of COVID-19, https://www.kff.org/coronavirus-covid-19/fact-sheet/three-years-of-covid-19/, 2023. (accessed 15 September 2024)

[2] World Health Organization, https://data.who.int/dashboards/covid19/cases. (accessed 15 September 2024)

[3] K.L. Andrejko, J.M. Pry, J.F. Myers, M. Mehrotra, K. Lamba, E. Lim, N. Fukui, J.L. DeGuzman, J. Openshaw, J. Watt, S. Jain, J.A. Lewnard, O. Covid-Case-Control Study Team, Waning of 2-Dose BNT162b2 and mRNA-1273 Vaccine Effectiveness Against Symptomatic SARS-CoV-2 Infection Accounting for Depletion-of-Susceptibles Bias, Am J Epidemiol 192 (2023) 895–907. 10.1093/aje/kwad017.

[4] B.J. Willett, J. Grove, O.A. MacLean, C. Wilkie, G. De Lorenzo, W. Furnon, D. Cantoni, S. Scott, N. Logan, S. Ashraf, M. Manali, A. Szemiel, V. Cowton, E. Vink, W.T. Harvey, C. Davis, P. Asamaphan, K. Smollett, L. Tong, R. Orton, J. Hughes, P. Holland, V. Silva, D.J. Pascall, K. Puxty, A. da Silva Filipe, G. Yebra, S. Shaaban, M.T.G. Holden, R.M. Pinto, R. Gunson, K. Templeton, P.R. Murcia, A.H. Patel, P. Klenerman, S. Dunachie, P. Consortium, C.-G.U. Consortium, J. Haughney, D.L. Robertson, M. Palmarini, S. Ray, E.C. Thomson, SARS-CoV-2 Omicron is an immune escape variant with an altered cell entry pathway, Nat Microbiol 7 (2022) 1161–1179. 10.1038/s41564-022-01143-7.

[5] Y. Oda, Y. Kumagai, M. Kanai, Y. Iwama, I. Okura, T. Minamida, Y. Yagi, T. Kurosawa, B. Greener, Y. Zhang, J.L. Walson, Immunogenicity and safety of a booster dose of a self-amplifying RNA COVID-19 vaccine (ARCT-154) versus BNT162b2 mRNA COVID-19 vaccine: a double-blind, multicentre, randomised, controlled, phase 3, non-inferiority trial, The Lancet infectious diseases 24 (2024) 351–360. 10.1016/S1473-3099(23)00650-3.

[6] Y. Oda, Y. Kumagai, M. Kanai, Y. Iwama, I. Okura, T. Minamida, Y. Yagi, T. Kurosawa, P. Chivukula, Y. Zhang, J.L. Walson, Persistence of immune responses of a self-amplifying RNA COVID-19 vaccine (ARCT-154) versus BNT162b2, The Lancet infectious diseases 24 (2024) 341–343. 10.1016/S1473-3099(24)00060-4.

[7] N.T. Ho, S.G. Hughes, V.T. Ta, L.T. Phan, Q. Do, T.V. Nguyen, A.T.V. Pham, M. Thi Ngoc Dang, L.V. Nguyen, Q.V. Trinh, H.N. Pham, M.V. Chu, T.T. Nguyen, Q.C. Luong, V.T. Tuong Le, T.V. Nguyen, L.T. Tran, A. Thi Van Luu, A.N. Nguyen, N.T. Nguyen, H.S. Vu, J.M. Edelman, S. Parker, B. Sullivan, S. Sullivan, Q. Ruan, B. Clemente, B. Luk, K. Lindert, D. Berdieva, K. Murphy, R. Sekulovich, B. Greener, I. Smolenov, P. Chivukula, V.T. Nguyen, X.H. Nguyen, Safety, immunogenicity and efficacy of the self-amplifying mRNA ARCT-154 COVID-19 vaccine: pooled phase 1, 2, 3a and 3b randomized, controlled trials, Nat Commun 15 (2024) 4081. 10.1038/s41467-024-47905-1.

[8] E. Dolgin, Self-copying RNA vaccine wins first full approval: what’s next?, Nature 624 (2023) 236–237. 10.1038/d41586-023-03859-w.

[9] M.C. Ballesteros-Briones, N. Silva-Pilipich, G. Herrador-Canete, L. Vanrell, C. Smerdou, A new generation of vaccines based on alphavirus self-amplifying RNA, Curr. Opin. Virol. 44 (2020) 145–153. 10.1016/j.coviro.2020.08.003.

[10] Y. Liu, Y. Li, Q. Hu, Advances in saRNA Vaccine Research against Emerging/Re-Emerging Viruses, Vaccines (Basel) 11 (2023). 10.3390/vaccines11071142.

[11] R. de Alwis, E.S. Gan, S. Chen, Y.S. Leong, H.C. Tan, S.L. Zhang, C. Yau, J.G.H. Low, S. Kalimuddin, D. Matsuda, E.C. Allen, P. Hartman, K.J. Park, M. Alayyoubi, H. Bhaskaran, A. Dukanovic, Y. Bao, B. Clemente, J. Vega, S. Roberts, J.A. Gonzalez, M. Sablad, R. Yelin, W. Taylor K. Tachikawa, S. Parker, P. Karmali, J. Davis, B.M. Sullivan, S.M. Sullivan, S.G. Hughes, P. Chivukula, E.E. Ooi, A single dose of self-transcribing and replicating RNA-based SARS-CoV-2 vaccine produces protective adaptive immunity in mice, Molecular therapy · the journal of the American Society of Gene Therapy 29 (2021) 1970–1983. 10.1016/j.ymthe.2021.04.001.

[12] T. Pepini, A.M. Pulichino, T. Carsillo, A.L. Carlson, F. Sari-Sarraf, K. Ramsauer, J.C. Debasitis, G. Maruggi, G.R. Otten, A.J. Geall, D. Yu, J.B. Ulmer, C. Iavarone, Induction of an IFN-Mediated Antiviral Response by a Self-Amplifying RNA Vaccine: Implications for Vaccine Design, J Immunol 198 (2017) 4012–4024. 10.4049/jimmunol.1601877.

[13] A.J. Geall, A. Verma, G.R. Otten, C.A. Shaw, A. Hekele, K. Banerjee, Y. Cu, C.W. Beard, L.A. Brito, T. Krucker, D.T. O’Hagan, M. Singh, P.W. Mason, N.M. Valiante, P.R. Dormitzer, S.W. Barnett, R. Rappuoli, J.B. Ulmer, C.W. Mandl, Nonviral delivery of self-amplifying RNA vaccines, Proceedings of the National Academy of Sciences of the United States of America 109 (2012) 14604–14609. 10.1073/pnas.1209367109.

[14] K.J. Hassett, I.L. Rajlic, K. Bahl, R. White, K. Cowens, E. Jacquinet, K.E. Burke, mRNA vaccine trafficking and resulting protein expression after intramuscular administration, Mol Ther Nucleic Acids 35 (2024) 102083. 10.1016/j.omtn.2023.102083.

[15] M. Buckley, M. Arainga, L. Maiorino, I.S. Pires, B.J. Kim, K.K. Michaels, J. Dye, K. Qureshi, Y. Zhang, H. Mak, J.M. Steichen, W.R. Schief, F. Villinger, D.J. Irvine, Visualizing lipid nanoparticle trafficking for mRNA vaccine delivery in non-human primates, bioRxiv (2024). 10.1101/2024.06.21.600088.

